# Soluble tubulin is locally enriched at mitotic centrosomes in *C. elegans*

**DOI:** 10.1101/543066

**Authors:** Johannes Baumgart, Marcel Kirchner, Stefanie Redemann, Jeffrey Woodruff, Jean-Marc Verbavatz, Frank Jülicher, Anthony A. Hyman, Thomas Müller-Reichert, Jan Brugués

## Abstract

During mitosis, the centrosome expands its capacity to nucleate microtubules. Understanding the mechanisms of centrosomal microtubule nucleation is, however, constrained by a lack of knowledge of the amount of soluble and polymer tubulin at mitotic centrosomes. Here we combined light microscopy and serial-section electron tomography to measure the amount of dimer and polymer at mitotic centrosomes in early *C. elegans* embryos. We show that a *C. elegans* one-cell stage centrosome at metaphase contains more than ten thousand microtubules with a total polymer concentration of 230 µM. Centrosomes concentrate soluble α/β tubulin by about tenfold over the cytoplasm, reaching peak values of 470 µM, giving a combined total monomer and polymer tubulin concentration at centrosomes of up to 660 µM. These findings support *in vitro* data suggesting that microtubule nucleation in *C. elegans* centrosomes is driven in part by concentrating soluble tubulin.

## Introduction

During mitosis, the pericentriolar material of centrosomes grows and increases their nucleation capacity many fold. This process, called centrosome maturation is thought to be essential for forming a mitotic spindle in many animal systems (Fu and Glover, 2012; Lawo et al., 2012; Mennella et al., 2014; Sonnen et al., 2012). Despite the importance of centrosome maturation in spindle formation, the mechanisms by which centrosomes increase their nucleation capacity are unknown. In *C. elegans* it is thought that increase of centrosome size is driven by accumulation of a coiled-coil protein called SPD-5, which forms the centrosome scaffold (Hamill et al., 2002). Growth of the scaffold is stimulated by the protein SPD-2 (Pelletier et al., 2004) and Polo kinase (Decker et al., 2011; Wueseke et al., 2016). SPD2/PLK1 dependent formation of the SPD-5 scaffold has been reconstituted *in vitro* (Woodruff et al., 2017). Similar pathways are thought to operate in *Drosophila* (Conduit et al., 2014). Once formed, it has been proposed that client proteins such as γ-tubulin, microtubules polymerases, and microtubule depolymerases, partition into the scaffold where they favor microtubule growth and nucleation (Woodruff et al., 2017). However, genetic evidence for the role of these microtubule-associated proteins in driving microtubule nucleation in mitosis is limited. In *C. elegans*, for instance, mutations or depletion of γ-tubulin only have marginal effects on nucleation (Hannak et al., 2002; O'Toole et al., 2012; Strome et al., 2001).

Recently, it has been suggested that nucleation could be driven in part by the partitioning of tubulin dimers into the pericentriolar material (PCM) (Woodruff et al., 2017). This would raise the tubulin concentration above the critical concentration for nucleation, thus driving microtubule formation. Supporting evidence for this idea comes from biochemical reconstitutions, which have shown that tubulin and other proteins can partition into reconstituted PCM and liquid drops of the microtubule associated protein tau (Hernandez-Vega et al., 2017; Woodruff et al., 2017), as well as experiments on the role of BugZ in assembling *Xenopus laevis* spindles (Huang et al., 2018). However, evidence for such ideas *in vivo* is currently lacking. In particular, we lack measurements for tubulin concentration of polymerized and unpolymerized tubulin at centrosomes, and the extent of local enrichment of tubulin within the living cell.

In this study, we quantitatively measured *in vivo* how α/βtubulin, in the form of soluble monomers as well as microtubule polymers, is distributed across the centrosome in *C. elegans* embryos. We show that centrosomes concentrate soluble α/β tubulin by about tenfold over the cytoplasm, reaching up to 470 µM. Based on our observations we propose that microtubule nucleation in mitotic *C. elegans* centrosomes is mediated in part by enriching tubulin locally.

## Results and discussion

To qualitatively measure the concentration of tubulin dimers at centrosomes, independent of its monomeric or polymeric state, we carried out confocal live-cell imaging of one-cell *C. elegans* embryos expressing GFP-tagged β-tubulin (GFP::TBB-2). The fluorescence intensity was roughly symmetric with respect to the centrosome center. To obtain the intensity profile along the radial direction, we used the centriole center as the origin of our analysis and averaged the signal along the circumferential direction in the region opposite to the spindle towards the cell cortex (Fig. 1A). The profiles of 19 embryos in metaphase showed a peak intensity of tubulin at a radial distance of around 1.0 µm, suggesting that tubulin is locally enriched at the outer shell of the centrosome (Fig. 1B). With further increasing distances we observed a monotonic decay to a plateau that extended away from the centrosome. This constant intensity outside the centrosome suggests a homogenous distribution in the cytoplasm of the total tubulin. At the very center of the centrosome, we detected a decrease of tubulin signal. However, the signal at the centrosome center was still higher compared to the plateau measured at the cortex. There was no significant difference in tubulin distribution between anterior and posterior centrosomes (Fig. 1B).

**Figure 1.**
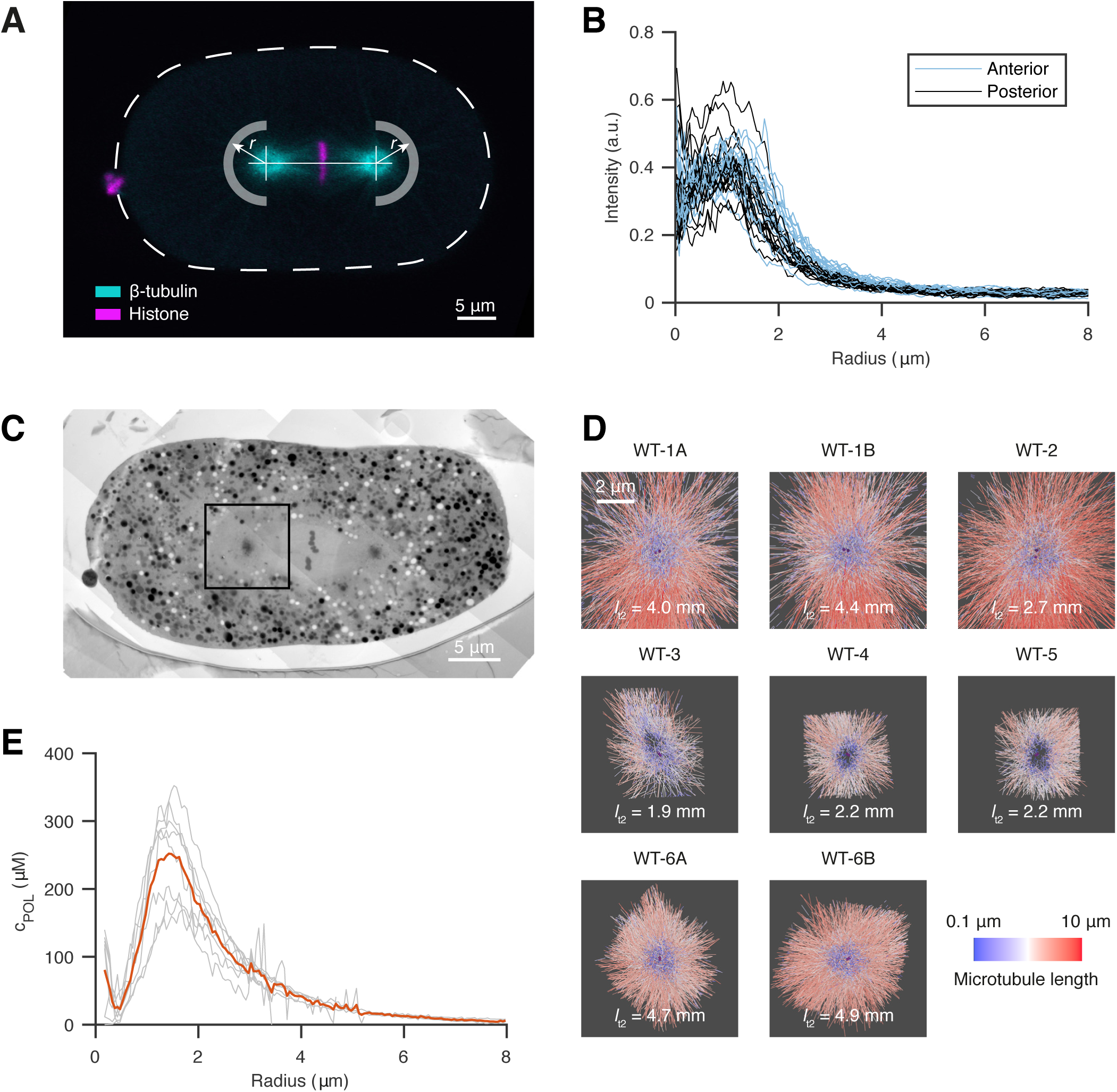
Light microscopy reveals a local enrichment of tubulin at the outer shell of the centrosome which co-localizes with a high density of microtubules recorded by electron tomography. (A) Confocal live-cell imaging of metaphase spindles in one-cell *C. elegans* embryos labeled with β-tubulin::GFP (cyan) and histone::mCherry (magenta). For the analysis, the centrosome centers were localized manually and fluorescent intensities of β-tubulin::GFP were extracted in radial distances as indicated by arrows for half planes away from the spindle and towards the cell cortex. The anterior side is orientated to the left. Scale bar, 5 µm. (B) Radial profiles of β-tubulin::GFP after subtraction of autofluorescence outside the cell (*n* = 19). There is no significant difference between anterior (light blue) and posterior centrosomes (dark blue). (C) Overview of a high-pressure frozen and serial sectioned one-cell embryo in metaphase. Scale bar, 5 µm. The black box indicates a representative area acquired for electron tomography. (D) Overview of all segmented centrosome models (*n* = 8). Six different embryos were used, A/B represent centrosomes of the same embryo. Microtubules are color coded by length in a logarithmical scale from short (blue) to long (red). *I*_t2_ represents the overall microtubule length up to a radius of 2 µm. (E) The segmented microtubules are analyzed in radial profiles as shown in A with respect to the local density as length per volume and are converted into the concentration of polymerized tubulin (c_POL_). For registration purposes, the center of the mother centriole is used to align the radial profiles.

To distinguish the soluble from the polymerized state, we performed serial-section electron tomography (EM) of eight centrosomes of six different cells in one-cell *C. elegans* embryos as previously reported (Fig. 1C-D) (Redemann et al., 2017) and extracted microtubule polymer concentration profiles (Fig. 1E). The number of microtubules at the centrosome of a one-cell stage embryo was more than ten thousand, with an average length of 1.1 µm. At the very center, a small peak was seen of about 100 µM due to the presence of the centriole microtubules, which we defined as the origin of our analysis. This peak was surrounded by a void region with a dip at 430 nm, comparable to the size of the dense interphase PCM layer seen by super-resolution light microscopy in *Drosophila* S2 cells and in human tissue culture cells (Lawo et al., 2012; Mennella et al., 2014). The microtubule density then increased up to a radial distance of about 1.4 µm, with the highest polymerized tubulin concentration in the range from 170 to 350 µM. This region coincided roughly with the outer edge of the PCM. The amount of polymeric tubulin then decayed monotonically with increasing distance from the centrosome.

To calibrate the measurement from light microscopy (LM), where both soluble and polymerized tubulin are labeled, we combined those measurements with the spatial and quantitative information from EM. Combining the quantitative EM data with the LM data required two compensatory procedures. First, correction for shrinkage during sample preparation for EM, and second, blurring of intensity profiles from LM due to the point-spread function (PSF) of the microscope. After accounting for these effects (see materials and methods), we were able to compare the radial concentration profiles from the two approaches. By calculating the difference between the calibrated intensity profile from light microscopy (c_TOT_) and the profile of the electron microscopy data (c_POL_) we obtained the concentration profile of soluble tubulin c_SOL_ = c_TOT_ – c_POL_ (Fig. 2A). This analysis revealed that the soluble tubulin concentration profile (c_SOL_) is significantly higher in the region of the centrosome. These analyses also showed that the total tubulin concentration profile had a maximum concentration of about 660 µM at radial distance of 1.0 µm from the centriole. Of this, 200 µM is polymerized and 460 µM is soluble tubulin.

**Figure 2.**
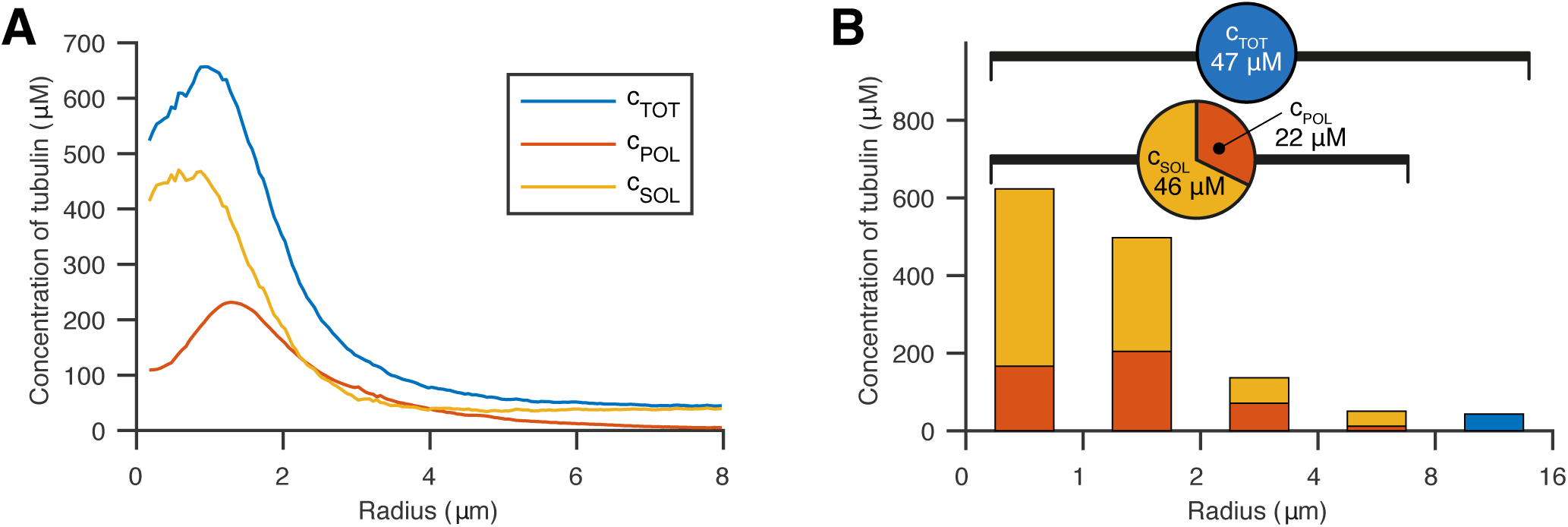
Calibrated intensity profiles from light microscopy by electron tomography data show the distribution of soluble and polymerized tubulin. (A) Concentrations of soluble (yellow), polymerized (red) and total (blue) tubulin concentration along the spindle axis after calibration. The soluble tubulin concentration is enriched at the centrosome and shows a peak concentration of about 405 µM at *r* = 0.8 µm. (B) Fractions of polymerized and soluble tubulin along the radial distance up to the data range of the EM reconstructions and the total concentration in the remaining part. The embryo is loaded with a total tubulin concentration of about 47 µM. Up to 8 µm, the overall tubulin is on average 68 µM of which 22 µM is polymerized.

We used these data to measure the total polymer and monomer concentration in the embryo. For the entire cell, we obtained a concentration of 47 µM, which is comparable to data from mass spectrometry (Saha et al., 2016). By integrating from the spindle axis up to a distance of 8 µm (the maximum distance at which we obtained EM data) we measured that about 32 % of tubulin was polymerized (about 22 µM), and the rest remained in the soluble state (about 46 µM) (Fig. 2B). Because the larger fraction of this recruited tubulin is freely available in the soluble state and only a smaller fraction is assembled into microtubules, this suggests that tubulin is not a limiting component in the process of microtubule nucleation at the centrosome. It also confirms that there must be mechanisms to concentrate tubulin at centrosomes, or the tubulin would simply diffuse out of the centrosome down the concentration gradient back into the cytoplasm.

To further explore the amount of soluble versus polymerized tubulin at mitotic centrosomes, we treated embryos with nocodazole to depolymerize microtubules (Fig. 3A) (Carvalho et al., 2011; Hannak et al., 2002; Strome et al., 2001). Electron tomography confirmed that there were only a few polymerized microtubules remaining at centrosomes (the total length of microtubules within a radius of 2 µm, covering the centrosome size, measures only about 4 % compared to the wild-type data, Fig. 3B). The concentration profile of soluble tubulin after nocodazole treatment showed a similar shape as that seen in wild-type embryos, with a peak of tubulin concentration close to the centrioles and a rapid decay outside of the centrosome (Fig. 3C). In contrast to wild type, the concentration profile showed no dip at the center, and peaked very close to the centrioles, suggesting that this dip is driven by microtubule polymerization. Supporting this idea, our tomographic data revealed that short microtubules are predominantly found at the centrosome center, whereas long microtubules tend to be located further outside (Figs. 1D, 4A-B). In addition, we found that microtubule polymerization velocity as measured with EB2 is lower closer to the centrosomes (Fig. 4C). We assume that the location of very short microtubules marks the region where nucleation predominantly takes place, and longer microtubules are in some way transported out to the periphery or stabilized (Fig. 4D).

**Figure 3.**
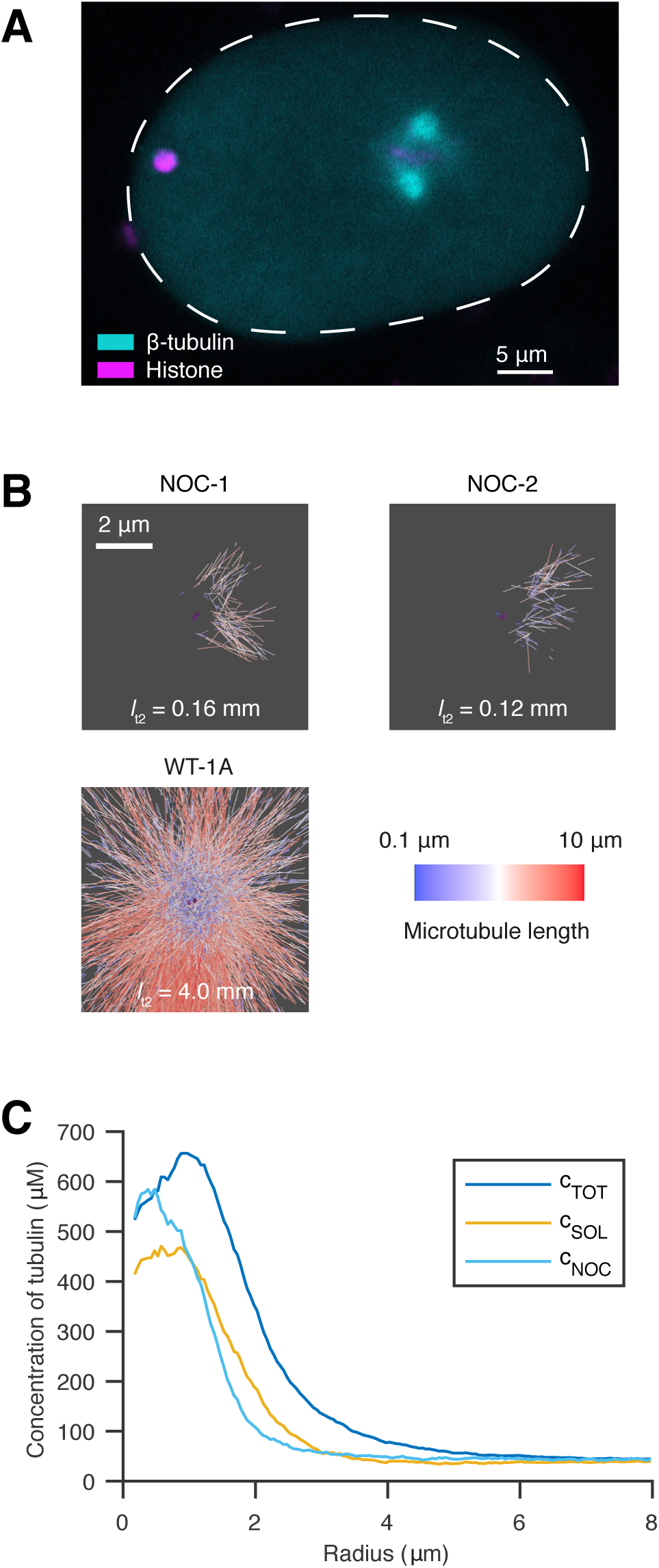
Effect of microtubule depolymerization on the centrosomal concentration of free tubulin. (A) Confocal live-cell imaging of metaphase spindles in one-cell *C. elegans* embryo labeled with β-tubulin::GFP (cyan) and histone::mCherry (magenta) 100 s after nocodazole treatment. The anterior side is orientated to the left. Scale bar, 5 µm. (B) Segmentations of microtubules at the centrosomes of nocodazole-treated embryos (*n* = 2, top, NOC) show a significantly reduced number of microtubules compared to wild-type embryos (bottom, WT). Microtubules are color-coded according to their length (short, blue; long, red). *I*_t2_ represents the overall microtubule length up to a radius of 2 µm. Scale bar, 2 µm. (C) Radial profiles of soluble tubulin after nocodazole treatment (light blue) compared to wild-type (soluble tubulin, yellow and total tubulin, blue).

**Figure 4.**
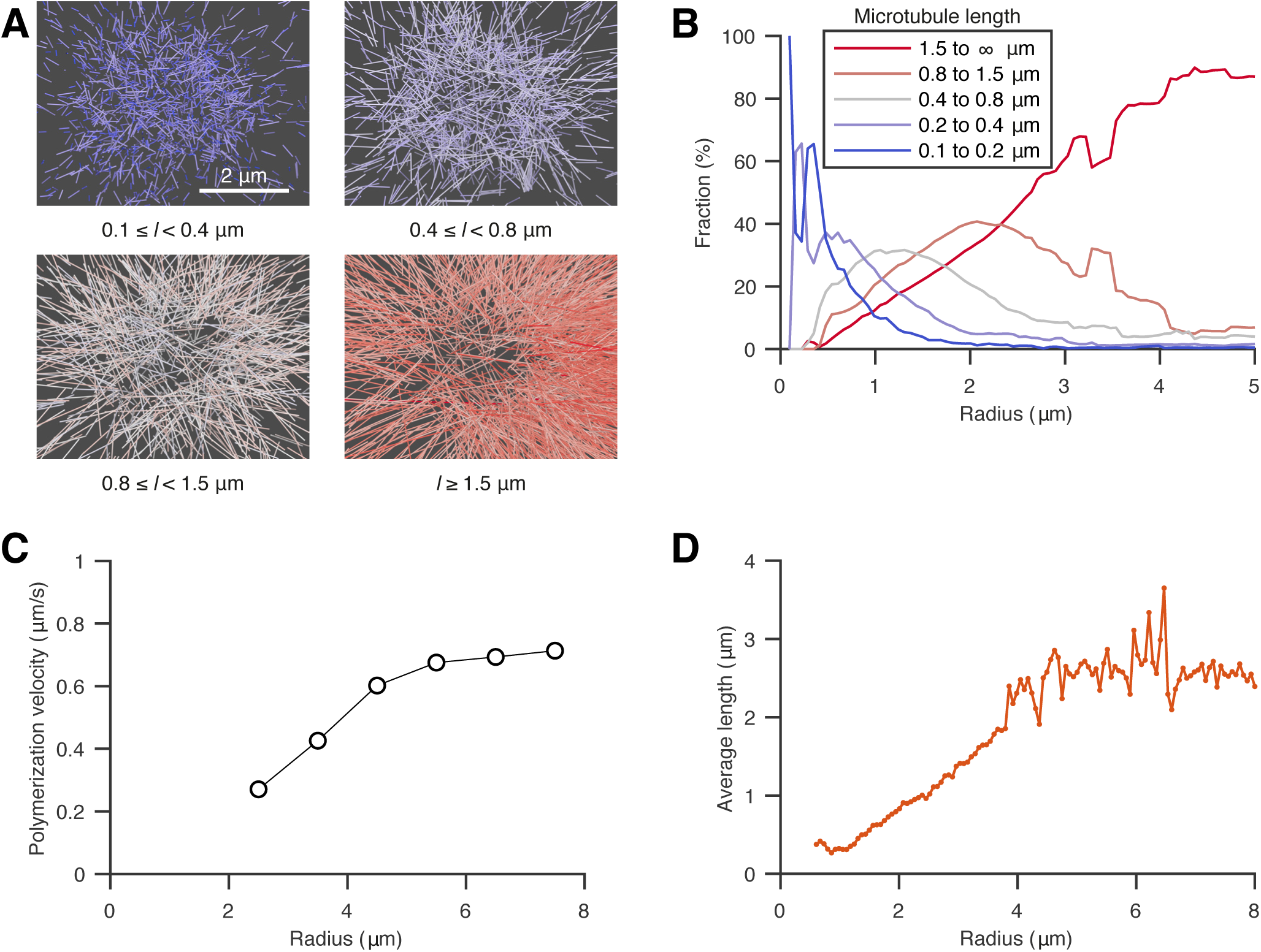
Localization of short microtubules at the centrosome. (A) Spatial graphs of a representative centrosome (WT-1A) showing groups of microtubules filtered by length from short (0.1 – 0.4 µm, blue) to long (> 1.5 µm, red). Very short microtubules cover predominantly the core of centrosomes, longer microtubules are found on the outer edge. Scale bar, 2 µm. (B) Plot of the fraction of microtubules of different length groups (short - blue to long - red) at specific radii from the centrosome center. Short microtubules are predominantly found at the centrosome, whereas longer microtubules tend to be located outside the centrosome. The radial coordinate is corrected for shrinkage (see materials and methods) (C) Spatial dependence of polymerization velocity measured from EB2 as a function of radial distance from the centrosome. (D) Average microtubule length as a function of the radial distance from the centrosome.

The mechanisms driving microtubule nucleation at mitotic centrosomes have long been a mystery. Although nucleation at interphase centrosomes is prevented by lowering the concentration of γ-tubulin by RNA interference (Hannak et al., 2002; O'Toole et al., 2012), similar treatments in mitosis have little effect on microtubule nucleation. The data presented in this paper show that centrosomes contain a considerable quantity of unpolymerized tubulin. This tubulin is constantly exchanging with the cytoplasm, suggesting that it can diffuse in the centrosome and drive microtubule growth. The typical critical concentration for spontaneous microtubule nucleation for bovine tubulin is thought to be 21 µM (Wieczorek et al., 2015). Therefore, the presence of such high concentrations of soluble tubulin supports the idea that increased concentration of tubulin could be a driving force for nucleation of microtubules at centrosomes.

We do not understand the mechanisms by which tubulin concentrates at centrosomes. However, it must do so in a form in which it can freely diffuse so that it can drive microtubule growth. One formalism would be to think of tubulin partitioning into the PCM, and that the tubulin concentration at the centrosome is defined by a partition coefficient (Woodruff et al., 2017). *In vitro* reconstitution experiments show that tubulin is not concentrated by the main components of the PCM, SPD-5 and SPD-2. Rather, by two microtubule associated proteins, ZYG-9, which is the *C. elegans* homologue of XMAP215, and TPXL-1, which is the *C. elegans* homologue of TPX2 (Ozlu et al., 2005). Interestingly, *in vitro* mutants in ZYG-9 that prevent tubulin binding, reduce tubulin association with centrosomes. This suggests that ZYG-9 could in part bind to and concentrate tubulin at centrosomes. However, *in vivo*, mutants in ZYG-9 do not prevent microtubule nucleation, indicating that in embryos, the process is more complicated and uses a combination of different tubulin binding proteins. Subsequent work combining electron tomography, light microscopy, and genetics, will be required to elucidate these mechanisms.

## Materials & methods

### Worm cultivation

The following strains were used in this study: wild-type N2 Bristol for electron tomography of WT-1A, WT-1B, and WT-2. MAS91 {unc-119(ed3) III; ItIs37[pAA64; pie-1::mCherry::HIS58]; ruIs57[pie-1::GFP::tbb + unc-119(+)]} for the other sets of electron tomography, light microscopic analysis and nocodazole treatment. TH315 {unc-119(ed3) III; ddEx23 [pie-1::GFP::SAS-4 (genomic introns, CAI 0.3); unc-119(+)]} for obtaining the point spread function at the imaging system for calibration. All strains were cultured on OP50 seeded NGM agar plates at 20 °C (Brenner, 1974). To enhance expression of fluorescent markers, worms were shifted to 25 °C 24 hours prior to light microscopic imaging.

### Light microscopy

Young hermaphrodites of the *C. elegans* MAS91 strain were dissected with syringe needles in M9 buffer on a coverslip (24 mm x 60 mm) to release the embryos. We took one-cell *C. elegans* embryos in metaphase, a stage where spindle growth is mainly completed and the microtubules are not exposed to high pulling forces. Imaging was performed using a Zeiss LSM 710 NLO multiphoton laser scanning microscope in single photon mode equipped with a Zeiss LCI Plan Neofluar 63x 1.3 NA water-immersion objective. Detection of the emitted fluorescent signal was carried out with a QUASAR detector with 32 channels and a dichromatic 488/594 nm beam splitter for excitation/emission splitting and subsequent linear unmixing of both acquired channels. A spectral prism slider was used for selecting the range of emission detection. Confocal stacks were acquired every 20 seconds covering a *z*-range of around 15 µm with a spacing of 0.388 µm. In total, we recorded stacks of 19 embryos in metaphase for wild-type analysis. The time point of metaphase was defined as the stack prior to anaphase onset in which clear segregation of chromosomes could be observed. Point spread function was measured using the fluorescent signal of SAS-4::GFP as reference beads inside the *C. elegans* embryo. For this, the strain TH315 was imaged under similar conditions. Individual SAS-4 spots (*n* = 21) from two- to sixteen-cell embryos were isolated and analyzed in cylindrical coordinates using axis-symmetry. Resulting in an averaged Gaussian shape intensity profile with a full-width half maximum of 0.423 ± 0.048 µm (mean ± STD) in the plane normal to the optical axis and 1.684 ± 0.336 µm (mean ± STD) along the axis (Supplementary Fig. 1A).

### Sample preparation for electron microscopy

Isolated early embryos were transferred into cellulose capillary tubes with a diameter of 200 µm (Leica Microsystems, Vienna, Austria) for high-pressure freezing. Embryos were observed with a stereomicroscope until metaphase and then high-pressure frozen using an EMPACT2 with a rapid transfer system (RTS, Leica Microsystems, Vienna, Austria) (Pelletier et al., 2006; Redemann et al., 2017). The following freeze substitution was carried out for 3 days at −90 °C using 1 % OsO_4_ and 0.1 % uranyl acetate using an automatic freeze substitution machine (EM AFS, Leica Microsystems, Vienna, Austria). Samples were embedded in a thin-layer of Epon/Araldite and polymerized at 60 °C for 3 days. Serial semi-thick sections (250-300 nm) were cut with an Ultracut UCT Microtome (Leica Microsystems, Vienna, Austria) and collected on Formvar-coated copper slot grids. Post-staining was performed with 2 % uranyl acetate in 70 % methanol and 0.4 % Reynolds lead citrate (Muller-Reichert et al., 2007). In this work, we only used the *C. elegans* MAS91 strain for calibration and analysis which exhibits a β-tubulin::GFP label.

### Electron tomography, 3D reconstruction and automatic segmentation of microtubules

Stained serial semi-thick sections were coated with colloidal gold (15 nm, Sigma-Aldrich) serving as fiducial markers for subsequent tomographic reconstruction. Dual-axis electron tomography was carried out using a TECNAI F30 TEM (FEI Company, Eindhoven, Netherlands) operated at 300 kV. Tilt series were acquired at every 1° in a range of ± 60° with a pixel size of 2.3 nm using a Gatan US1000 CCD camera (2k x 2k). For tomographic reconstruction, the IMOD software package (http://bio3d.colourado.edu/imod) was used (Kremer et al., 1996). With this software, the computation is based on an R-weighted back-projection algorithm for each tilt axis (Gilbert, 1972). After reconstruction, tomograms were flattened and trimmed to remove unsubstantial outer edges of the volume with no information. For segmentation and automatic tracing of microtubules, we used the AMIRA software with an extension to the filament editor (Weber et al., 2012). After automatic segmentation of microtubules, three-dimensional models were visually inspected and manually corrected. This correction included manual tracing of undetected microtubules, extending and combining individual traces as well as removing incorrectly traced structures, such as membranes and other cellular components. Using the Amira software, corrected three-dimensional models were then stitched in *z* to obtain complete volumes of the recorded centrosomes (Weber et al., 2014). In total, we recorded eight wild-type and two nocodazole-treated centrosomes covering an average range of 5 µm in the slice plane. Merged data sets consist of at least seven subsequent sections for the wild-type preparations and four for the nocodazole-treated embryos.

### Calibration of light and electron microscopy

Light microscopy provides a qualitative measure of local tubulin concentration. There is no distinction in signal intensity between tubulin polymerized into microtubules and in solution. We corrected for the background signal by subtracting the median value outside the cell. Also, we controlled for the auto-fluorescence signal by measuring intensity values in unlabeled N2 wild-type embryos (*n* = 26). This signal was always below 10 % of the measured intensity and decayed towards the centrosome. The coordinates of the centrosome centers, serving as the origin of our radial profiles, were manually defined by observing the data sets in FIJI in orthogonal views. Radial profiles were extracted in the image plane of the stack, which contained the manually defined center of the centrosome. For each radial position, all values in the circumferential direction were averaged for the half plane away from the spindle. Each radial profile was normalized with the integrand of the respective intensity profile from the center to *r* = 8 µm (the maximum distance at which we obtained EM data).

Electron tomography provides a quantitative measure of polymerized tubulin as only assembled microtubules are detectable. In total, we segmented the microtubules of 8 wild-type one-cell *C. elegans* metaphase centrosomes. For registration purposes, we used the center of the mother centriole as the origin of analysis. The spindle axis was estimated either by the coordinates of the opposite mother centriole or if not available by manual inspection of the data set. The segmented microtubules were analyzed with respect to the microtubule density by computing locally the microtubule length per volume. Assuming all microtubules would have the same direction and cover the entire volume, this corresponds to the number of microtubules per area. Microtubules in the mitotic spindle have predominantly 11 protofilaments (Chaaban et al., 2018), which corresponds to about 1360 tubulin dimers per micrometer; the corresponding length per volume relates to the molar concentration of polymerized tubulin into microtubules as c/(µM) = 2.255 ρ/(µm/µm^3^). We estimated the density on a three-dimensional grid with an equidistant spacing of 100 nm. We took the boundary of the data sets into account and averaged the density values in a spherical coordinate system in the circumferential direction with respect to the spindle axis.

To compare the data from electron with those from light microscopy we had to correct artifacts of the respective methods. To consistently calibrate the light with the electron microscopy signal and combine both radial profiles, we compensated the sample shrinkage during the workflow of electron tomography. For this purpose, we manually measured the dimensions of embryo length and width in the maximum projection of confocal stacks before preparation for electron tomography and in serial sections after preparation (*n* = 9) and estimated a shrinkage factor of about 27.5 %. One possibility to correct for the point-spread function is to deconvolve the raw light microscopy data. We discarded this method, however, as the PSF is not precisely known and the data are rather noisy. Instead, we convolved the shrinkage-corrected EM data set with the PSF to compare profiles with the same recording conditions. For the convolution, we used the three-dimensional EM data and convolved them with the Gaussian PSF on a slice that encloses the spindle axis.

To calibrate the light microscopy data we used the assumption that soluble tubulin is homogenously distributed outside of the centrosome (*r* > 4 µm) and that the spatial variation there is solely due to the spatial changes of the concentration of polymerized tubulin. To implement this, we calibrated the signal of the light microscopy by minimizing the variance of the computed soluble tubulin in the region from 4 to 8 µm in three reconstructions. The profiles of polymerized tubulin in five additional datasets have a similar profile up to about 4 µm.

### Averaged concentrations

For the averages of total and polymerized tubulin we used the calibrated profiles and integrated them numerically for the radial ranges as specified. We used spherical coordinates for the two half-spheres away from the spindle and cylindrical coordinates for the region between the centrioles. In both cases we used averaged profiles depending only on the distance to the center of the centrosome, either the radius or the position on the spindle axis with respect to the nearby centrosome. For radii larger than 8 µm we had only data from light microscopy. Consequently, we report only total concentrations for this region and the entire cell.

## Drug treatment

For the nocodazole experiment, worms were treated with *perm-1 (RNAi)* feeding (T01H3.4) for 17 hours at 25 °C to permeabilize the eggshell (Timmons and Fire, 1998). Worms were dissected in a mixture of *perm-1* and M9 buffer (1:1.5) and embryos were selected and transferred into a microdevice as described in (Carvalho et al., 2011). For microtubule depolymerization, the medium within the microdevice was replaced at the desired stage of nuclear envelope breakdown with fresh medium containing a final concentration of 50 µg/ml nocodazole (Sigma, #M1404). Imaging could be carried out continuously and the effect of depolymerization was visible 1 min after drug addition. The light microscopic acquisition was carried out with the same settings as for the wild-type experiment. This allowed us to use the same calibration coefficient as for the wild-type experiment.

### Data analysis

Data analysis was carried out using either the AMIRA software (Zuse Institute Berlin, Germany) or Matlab (R2017b, The MathWorks, Natick). To reduce a bias resulting from errors in the tracing algorithm, microtubules shorter than 100 nm were excluded from all analyses. The microtubules are very stiff and consequently close to straight lines. For simplicity, we treated them as straight lines to compute the local density. For the fractions of microtubules based on length, microtubules were grouped by the end-to-end length. At a given radial position the number of microtubules in each group was counted and compared to the total number crossing this radius.

The semi-thick sections cut microtubules at their boundary. For an unbiased estimate of the local average length, a density profile was reconstructed by convolving the density of the minus ends, which are proximal to the centrosome, with an exponential length distribution. The local average length of this distribution (Fig. 4D) was set to reproduce the actual density profile (Fig. 1E).

There are two distinct experimental methods involved with light and electron microscopy to obtain the profile of soluble tubulin. There are many possible sources of errors, and not all can be quantified here. However, the lower bound calibration (which sets the minimum of the soluble tubulin to zero, Supplementary Fig. 1B) shows that even in this situation soluble tubulin is enriched ten-fold at the poles, similar to our variance calibration. The overall density is 59 % smaller than the estimate based on the variance approach and the qualitative shape of the profiles is maintained, except a minimum for the soluble tubulin.

## Acknowledgements

The authors would like to thank Norbert Lindow, Steffen Prohaska, Martin Wigert and Oliver Wüseke and the members of the LM and EM facility at MPI-CBG for discussions and technical assistance. S. Redemann received funding from the Faculty of Medicine Carl Gustav Carus of the TU Dresden (Frauenhabilitationsstipendium). Research in the Hyman and Jülicher groups is supported by the European Comission’s 7th framework Programme grant Systems Biology of Mitosis (FP7_HEALTH-2009-241548/MitoSys). Research in the Müller-Reichert group is supported by funds from the Deutsche Forschungsgemeinschaft (MU 1423/8-1 and 8-2). Research in the Brugués group is funded by the Human Frontiers Science Program (CDA00074/2014). Research in the Jülicher, Hyman, and Brugués groups is funded by the Deutsche Forschungsgemeinschaft (DFG, German Research Foundation) under Germany’s Excellence Strategy – EXC-2068 – 390729961– Cluster of Excellence Physics of Life of TU Dresden.

## Contributions

This work represents a truly collaborative effort. Each author has contributed significantly to the findings and regular group discussions guided the development of the ideas presented here.

---

**figure S1.**
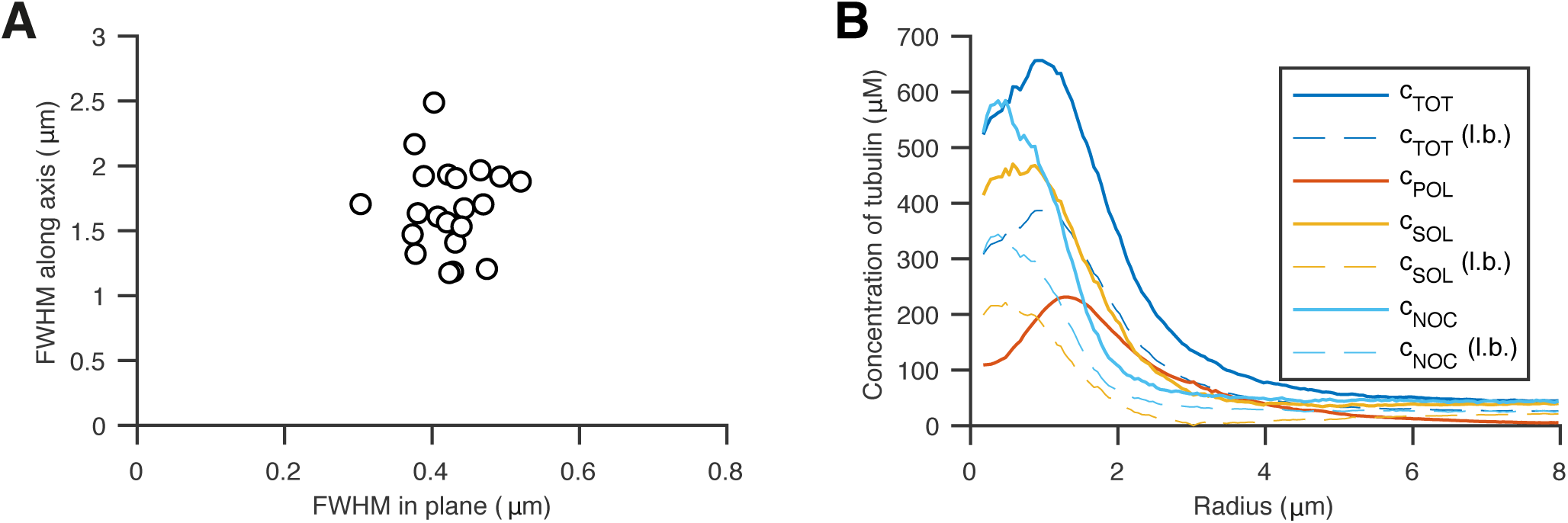
Size of point spread function and lower bound of concentration profiles. (A) Individual SAS-4 spots (*n* = 21) recorded in embryos were fitted with an axis-symmetric gaussian intensity profile. Plot of the full-width at half maximum (FWHM) values in the plane normal to the optical axis with respect to along the axis. (B) Density curves corresponding to figures 2A and 3C (solid lines) replotted together with the lower bound calibration of the light microscopy data (l.b., dashed lines). The lower bound of the calibration coefficient for the light microcopy is obtained by imposing that all concentrations must be positive (see materials and methods).

